# Triple Negative Breast Cancer-derived Small Extracellular Vesicles as Modulator of Biomechanics in target cells

**DOI:** 10.1101/2022.02.28.481921

**Authors:** Beatrice Senigagliesi, Giuseppe Samperi, Nicola Cefarin, Luciana Gneo, Sara Petrosino, Mattia Apollonio, Federica Caponnetto, Riccardo Sgarra, Licio Collavin, Daniela Cesselli, Loredana Casalis, Pietro Parisse

## Abstract

Extracellular vesicle (EV) mediated communication has recently been proposed as one of the pivotal routes in the development of cancer metastasis. EVs are nano-sized vesicles swapped between cells, carrying a biologically active content that can promote tumor–induced immune suppression, metastasis and angiogenesis. Thus, EVs constitute a potential target in cancer therapy. However, their role in triggering the premetastatic niche and in tumor spreading is still unclear. Here, we focused on the EV ability to modulate the biomechanical properties of target cells, known to play a crucial role in metastatic spreading. To this purpose, we isolated and thoroughly characterized triple-negative breast cancer (TNBC)-derived small EVs. We then evaluated variations in the mechanical properties (cell stiffness, cytoskeleton/nuclear/morphology and Yap activity rearrangements) of non-metastatic breast cancer MCF7 cells upon EV treatment. Our results suggest that TNBC-derived small EVs are able to directly modify MCF7 cells by inducing a decrease in cell stiffness, rearrangements in cytoskeleton, focal adhesions and nuclear/cellular morphology, and an increase in Yap downstream gene expression. Testing the biomechanical response of cells after EV addition might represent a new functional assay in metastatic cancer framework that can be exploited for future application both in diagnosis and in therapy.

## 1 INTRODUCTION

Breast Cancer is the most frequently diagnosed malignancy and stands as the leading cause of cancer mortality in women worldwide (Momenimovahed & Salehiniya, 2019). Triple-negative breast cancer (TNBC), in particular, is the most aggressive breast cancer subtypes and with a poor prognosis due to the absence of targetable receptors, such as estrogen receptor (ER), progesterone receptor (PR) and epidermal growth factor receptor-2 (HER2), and to the high propensity for metastatic progression (Yeo, 2015). Nowadays, chemotherapy is the main treatment in both early and advanced stage of TNBC (Shang et al., 2018). Unfortunately, approximately 80% of TNBC patients show an incomplete response to conventional chemotherapy, disease recurrence, and metastasis formation after surgery (Jhan & Andrechek, 2017; Nakashoji et al., 2017). Therefore, shedding light on the biological mechanisms of the metastatic progression in TNBC is urgent to track down novel therapeutic approaches for effective interventions (Yao et al., 2021). The development of metastasis requires a series of stages that lead to the formation of secondary tumor sites in distant organs: metastatic cancer cells leave the primary site, pass through the basement membrane and extracellular matrix (invasion process), penetrate and survive in lymphatic or vascular circulation (intravasation), leave vessels (extravasation), and ultimately create pre-metastatic niche for the formation of the secondary tumor sites (Martin et al., 2014; Shibue & Weinberg, 2017). Nowadays, the capacity of cancer cells to undergo different phenotypic changes is well recognized.

The recent literature has provided evidences for a direct correlation between the metastatic potential of cancer cells and their biomechanical properties: deformable, softer cancer cells can migrate more easily through the narrow pores of the matrix and vessels boosting the processes of invasion, intravasation, and extravasation (Alibert et al., 2017; Lekka, 2016; Q. Luo et al., 2016). However, the specific nature of these mechanisms remains to be understood. In fact, cell biomechanical changes include complex transformations at the level of nucleus, cytoskeleton, and plasma membrane that are due to the mutual interaction with the extracellular microenvironment (Alibert et al., 2017; Lüchtefeld et al., 2020). On the other hand cell motility, to which the metastatic potential of cancer cells is directly related, depends on the bidirectional interplay between actin and microtubule organization and expression, and several cytoskeletal regulators (Dogterom & Koenderink, 2019; Etienne-Manneville, 2004). The cancer community has recently also emphasized the role of the tumor-secreted extracellular vesicle (EVs) in the regulation of tumor progression and metastasis (Bao et al., 2021; Becker et al., 2016). EVs are nano-sized particles delimited by a lipid bilayer, capable of transferring functional cargos (e.g. proteins, nucleic acids, and lipids) from donor to target cells, in which they activate combinatorial effects (Tschuschke et al., 2020). Since different tissues and organs throughout the body release EVs, analysis of EVs could give useful information in the early diagnosis, progression, and therapy monitoring of diseases (Boukouris & Mathivanan, 2015; Kalluri & LeBleu, 2020). Moreover, being released in the extracellular space, EVs can be detected non-invasively in body fluids (Boukouris & Mathivanan, 2015). According to the vast majority of the EV community, vesicles are divided into two big classes, “small EVs” (sEVs) and “medium/large EVs” (m/lEVs), with dimensions < 200 nm and > 200 nm, respectively (Thery et al., 2018). M/lEVs are mainly formed by the direct outward budding of the cellular plasma membrane (Cai et al., 2007), while sEVs have usually an endocytic origin (Bebelman et al., 2018). The involvement of EVs in tumor-tumor and tumor-stromal cell (non-malignant cells that surround the primary tumor) communication of both primary tumor progression and metastasis formation has been documented (Becker et al., 2016; Chen et al., 2017; Maacha et al., 2019). Concerning TNBC, it has been shown that TNBC cells can transfer oncogenic proteins, mRNAs, and miRNAs to target cells through EVs by promoting metastatic spreading and pre-metastatic niche formation (Green et al., 2015). Although EVs are known to be the main putative agents at the base of these processes (Ozawa et al., 2018; Peng et al., 2018), further attention is still needed to grasp the mechanisms through which they operate. Understanding the ways EVs deliver their cargo and affecting host cells properties is crucial to ultimately exploit them as novel therapeutic vehicles (Bahreyni et al., 2020). In particular, there is evidence that small extracellular vesicles (small EVs) are involved in modulation of cellular signaling pathways and metabolic state of target cells (Maia et al., 2018). Yet, to our knowledge, the role played by the small EVs in the modulation of recipient cell biomechanics has been poorly investigated. Here, we focused on the isolation and characterization of TNBC-derived small EVs and on the analysis of the phenotype changes they induce in non-metastatic target cells. We demonstrate for the first time a direct involvement of small EVs in the modulation of cytoskeleton, adhesion, nuclear/cellular morphology, and, as a consequence, on the biomechanical properties of the entire cells. Our findings might open new routes for the diagnosis and future therapy of this aggressive, target-orphan breast cancer subtype (Goh et al., 2020), as well as new insights into EV activity. Such findings could also be extended to other classes of EVs or to other diseases, where biomechanical properties play a crucial role (Lee & Lim, 2007).

## 2 MATERIALS AND METHODS

### 2.1 Small Extracellular Vesicles Isolation

#### 2.1.1 Cell cultures and experimental conditions for small extracellular vesicle isolation

MCF7 and MDA-MB-231 breast cancer cell lines were cultivated in DMEM (Dulbecco’s Modified Eagle’s Medium High Glucose with Sodium Pyruvate with L-Glutamine, EuroClone, ECM0728L) supplemented with 10% FBS (Fetal Bovine Serum South America origin EU, EuroClone, ECS0180L) and 1% Penicillin/Streptomicin (100X, EuroClone, ECB3001D). Cell lines were grown at 37°C in humidified 5% CO_2_ incubator and split every 2-3 days according to their confluence. The culture and harvesting conditions, such as passage number and seeding confluence, were maintained the same and regular checks for Mycoplasma contamination were performed on cells for vesicle isolation.

#### 2.1.2 Small Extracellular vesicle isolation by ultracentrifuge

MDA-MB-231 cells (2.5 × 10^6^) were grown in 175 cm^2^ flask in DMEM with 10% FBS for 2-3 days in order to avoid cellular stress; after that the cells were washed twice with PBS and, then, three times with DMEM without FBS to reduce the presence of serum contaminants (e.g. serum vesicles, albumin, RNA or proteins). After 24h in DMEM without FBS the small EVs were collected. The supernatant was centrifuged at 300 g for 10 minutes at 4°C to pellet cells and cell debris. The resulting supernatant was filtered using a 0.2 µm filter to remove the medium/large EVs, big circulating proteins and cell debris. The filtered supernatant was transferred into Amicon Ultra-15 centrifugal filters (Ultracel-PL PLHK, 100kDa cutoff, Merck Millipore, UFC9100) and centrifuged at 4,000 g for 40 minutes at 4°C, in order to concentrate the medium to use the ultracentrifuge tubes with a reduced volume capacity for the small EV isolation (8.9 ml polypropylene centrifuge tube, Beckman Coulter, 361623). The tubes were filled with PBS to reach the final volume and samples were ultracentrifuged at 120,000 g for 60 minutes at 4°C (70.1 Ti rotor, k-factor 36, Beckman Coulter, Brea, CA, USA). Finally, the supernatant was removed, the pellet resuspended in PBS, and small EVs were stored at -20**°**C for short term periods.

### 2.2 Small Extracellular Vesicles Characterization

#### 2.2.1 Scanning Electron Microscopy

Scanning Electron Microscopy (SEM) images were acquired with a Zeiss Supra40 SEM. Imaging was performed at low acceleration voltage (5 keV) by detecting secondary electrons. The silica slide was cleaned with acetone and isopropanol and a drop of Poly-L-Lysine (Sigma-Aldrich) was added, in order to facilitate the capture of small EVs via electrostatic interactions. Subsequently, the excess of Poly-L-Lysine was removed by performing two washes with H_2_O Milli-Q. Then, 10 µL of small EVs were spotted on the treated silica slide. The vesicles were mixed directly on the silica slide with an equal volume of 5% glutaraldehyde solution prepared in PBS to allow the vesicle fixation. The mixture was incubated for 30 minutes. The sample was washed and dehydrated with increasing ethanol solution, until it dried at room temperature. PBS was used as negative control. Before the scanning, sample was sputter-coated with a thin layer of Au/Pd (thickness of approximately 5 nm) to assure conductivity. For each sample, 10 different areas of ∼ 14 µm x 10 µm were imaged. SEM images were analyzed with Gwyddion^®^ software (Nečas & Klapetek, 2012). Vesicle diameters were obtained by applying a threshold to the images and then evaluating the grain distributions.

#### 2.2.2 Atomic Force Microscopy

Atomic Force Microscopy (AFM) images were acquired using a commercially available microscope (MFP-3D Stand Alone AFM from Asylum Research, Santa Barbara, CA). Measurements were carried out at room temperature working in dynamic AC-mode. Commercially available silicon cantilevers (BL-AC40TS-C2, Olympus Micro Cantilevers, nominal spring constant 0.09 N m−1 and resonant frequency 110kHz) have been chosen for imaging in liquid. For the AFM imaging of the small EVs, a freshly cleaved muscovite mica sheet (Ruby Muscovite Mica Scratch Free Grade V-1, Nanoandmore GMBH, USA) was incubated with a drop of Poly-L-Lysine (Sigma-Aldrich) for 15 minutes at room temperature. Subsequently, the excess of Poly-L-Lysine was removed by performing two washes with H_2_O Milli-Q. A drop of small EV suspension was loaded to the poly-lysine-coated mica surface at room temperature for 15 minutes to allow vesicles to bind the surface via electrostatic interactions. The PBS was used as negative control. For each sample, 5 images with 10 µm x 10 µm of scan size and with a resolution of 1024 × 1024 pixels (pixel size ∼ 10 nm x 10 nm) were acquired. The AFM images were analyzed with the Gwyddion^®^ software. Vesicle heights and diameters were obtained by applying to the images a threshold of 10 nm in height and evaluating, then, the grain distributions of all grains higher than 10 nm.

#### 2.2.3 Nanoparticle Tracking Analysis

Concentration and particle size distribution of purified small EVs derived from MDA-MB-231 were obtained by Nanosight (LM10, Malvern system Ltd., U.K.), equipped with a 405 nm laser. Each sample, once properly diluted, was recorded for 60 seconds with a detection threshold set at maximum. Temperature was monitored throughout the measurements. Vesicle size distribution and an estimated concentration of NTA (Nanoparticle Tracking Analysis) profiles were obtained from the given raw data files.

#### 2.2.4 Western blot

MDA-MB-231 cells and 231_sEVs were dissolved in SDS sample buffer (125 mM Tris/HCl pH 6.8, 4% w/v SDS, 20% glycerol, traces of bromophenol blue and 0.2 M DTT) and heated for 5 minutes at 96 °C (except samples for recognition of tetraspanins). Proteins were separated by SDS-polyacrylamide gel electrophoresis and transferred at 4 °C for 16 hours to a nitrocellulose membrane (Ø 0.2 µm GE Healthcare, Whatman, 10401396) through a wet transfer system (transfer buffer: 20% methanol, 25 mM Tris, 200 mM Glycine). Membrane was stained with Red Ponceau solution (0.2% Red Ponceau S, 3% trichloroacetic acid, 3% sulfosalicylic acid), incubated in agitation for 10 minutes. After blocking the membranes (5% NFDM – non-fat dry milk (w/v) and 0.1% (v/v) Tween 20 in PBS), blots were incubated for 1 hour at RT with primary antibodies. Incubation with horseradish peroxidase-conjugated secondary antibodies for 1 hour was performed after three washes of the membranes with blocking solution. After three final washes with blocking solution and two with PBS, chemiluminescence substrate ECL kit (Thermo Scientific, 2106) was added to the membrane in order to visualize the target proteins through Bio-Rad ChemiDoc™ Imagers. The primary antibodies used were: anti-CD63 (1:50, in native conditions, Santa Cruz Biotechnology, sc-5275), anti-Tsg 101 (1:50, Santa Cruz Biotechnology, sc-7964), anti-Calnexin (1:80, Santa Cruz Biotechnology, sc-23954), and anti-Albumin (1:50, Santa Cruz Biotechnology, sc-374670).

### 2.3 Functional Experiment: Small Extracellular Vesicle uptake in target cell

#### 2.3.1 Bradford assay

Protein concentration via Bradford assay was used to obtain an estimation of the quantity of small EVs for downstream application. For sample preparation, small EVs and cells were lysed in RIPA buffer in order to extract proteins. Solutions were centrifuged at 14,000 g for 10 minutes, recovering the supernatant, in order to eliminate cell debris. A small volume of samples (different BSA solutions and the unknown sample) was deposited on a 96-well plate. Afterwards, 200 µL of Coomassie Brilliant Blue G-250 (Bradford-Solution for protein determination, EuroClone, APA69320500) was added to each sample. Incubation for 10 minutes led to a stable protein–dye complex that was monitored at 595 nm using a spectrophotometer TECAN infinite F200 PRO (Tecan Trading AG, Switzerland). The vesicle protein amount was calculated using a BSA calibration curve in a range of 0.05–2 mg/mL. Each sample was analyzed in triplicate (in three different wells).

#### 2.3.2 Proliferation assay

MCF7 and MDA-MB-231 cells (1 × 10^5^ for 24h and 0.5 × 10^5^ cells for 48h of incubation time) were seeded in a 24-well plate and were left to grow for 24h. Afterwards, cells were washed and fresh culture medium containing small EVs derived from MDA-MB-231 at different concentrations (0.5 μg/μL, 0.1 μg/μL and 0.2 μg/μL) was added. PBS was used as negative control. Each experimental condition was analyzed in triplicate (in three different wells). MCF7 cells were left to incubate with vesicles for 48h. Then, target cells were collected and counted.

#### 2.3.3 Immunofluorescence

Immunofluorescence images were carried out using a microscope (Inverted Research Microscope Eclipse Ti, Nikon) equipped with an epi-fluorescence illuminator or a 488 nm laser for Total Internal Reflection Fluorescence (TIRF) application. For sample preparation, cells were fixed in 4% paraformaldehyde for 20 minutes, washed in PBS, permeabilized with 0.5% PBS-TWEEN for 10 minutes and 0.1% PBS-TWEEN for 5 minutes (three times). Subsequently, cells were blocked in 1% BSA in 0.1% PBS-TWEEN for 60 minutes. Antigen recognition was performed by incubating primary antibody in a humidified chamber for different times (Phalloidin, Invitrogen, A12381; anti-Vinculin, Invitrogen, 42H89L44; anti-pFAK, Cell signaling, 3283S) and with anti-mouse/rabbit Alexa Fluor 488-594 (Invitrogen, A11008, A11005) as secondary antibody in a humidified chamber for 60 minutes. Nuclei were stained with DAPI (Sigma Aldrich). Images were analyzed by using ImageJ^®^.

#### 2.3.4 Force Spectroscopy Atomic Force Microscopy

Force spectroscopy analysis of cells was carried out by using a Smena AFM (NT-MDT Co., Moscow, Russia) mounted on an inverted fluorescence microscope (Nikon Eclipse Ti-U). A silicon spherical tip with a diameter of 20 µm (Tip: CSG01 cantilever from NT-MDT Smena, k = 0.006-0.012 N/m) was used, in order to collect the global stiffness of each cell. For sample preparation, cells were fixed in 4% paraformaldehyde for 20 minutes, washed in PBS and stained with DAPI. Cells were measured in PBS buffer with 1% penicillin/streptomycin at RT. Despite PFA fixation can induce alteration in the cell stiffness (Kim et al., 2017), it is known that the relative variations stiffness after treatments remain statistically significant even after fixation (Grimm et al., 2014). Moreover, the fixation avoids the damaging due to the cell aging and it is needed for the immunofluorescence studies. For each sample, at least 30 (or 60) randomly chosen cells were measured and analyzed. Force spectroscopy measurements were performed at constant speed (2 μm/s), with a maximum indentation of 0.5 μm and with a force applied to the sample of 1-2 nN. Elastic modulus values (E, kPa) were determined by fitting the obtained force-displacement curves with Hertz model by using AtomicJ^®^ software (Hermanowicz et al., 2014).

#### 2.3.5 Quantitative Real-Time Polymerase Chain Reactions

Quantitative Real-Time PCR (RT-qPCR) technique was used to quantify gene expression of Yap downstream genes (CTGF, CYR61, and ANKRD1) in cells. Cells were lysed by TRIzol and RNA was extracted using EuroGOLD TriFast reagent (Euroclone), according to manufacturer’s instructions. Purified RNA samples were quantified at Nanodrop Spectrophotometer device, by evaluating ng/μl concentration, protein and phenol/ethanol contaminations. The integrity of extract material was detected analyzing the coil and supercoil strips formation. For RNA expression analysis, 0.5 μg of total RNA sample (100 ng/μl) was retrotranscribed in stable cDNA with iScriptTM Advanced cDNA Synthesis Kit (Bio-Rad). Genes of interest were amplified with Itaq UniversSYBR Green (Bio-Rad), according to manufacturer’s instructions. A CFX ConnectTM Real-Time PCR System (Bio-Rad) was used to perform Real-Time PCR. All quantitative real-time PCR (qRT-PCR) results were normalized to histone H3.

#### 2.3.6 Data processing and statistics

The showed results are representative. At least 3 other independent experiments with the same trend were obtained. Data processing were performed by using Igor Pro^®^, Origin Graph^®^, and Microsoft Excel^®^ software. Significance of data differences was established via two-tailed Student’s t-test for immunofluorescence analyses and real-time PCR experiments and via one-way Anova for cell proliferation assay (* = *p* < 0.05; ** = *p* < 0.01; *** = *p* < 0.001; **** = *p* < 0.0001, respectively). Whereas, for force spectroscopy AFM analyses the non-normally distributed Young’s moduli (investigated via Shapiro-Wilk test) were compared by using the Wilcoxon test or Kruskal-Wallis one-way Anova test (* = *p* < 0.05, ** = *p* < 0.01, *** = *p* < 0.001 and **** = *p* < 0.0001, respectively).

## 3 RESULTS

TNBC-derived EVs have been shown to promote proliferation and drug resistance in non-tumorigenic breast cancer (Ozawa et al., 2018) and to induce an increase in cell migration proportional to the metastatic potential of donor cells (Harris et al., 2015). Moreover, small EVs derived from breast cancer-associated stromal cells have been found to promote proliferation and migration via Hippo signaling pathway activation in non-invasive breast cancer cells (Nardone et al., 2017). Considering all this evidence, we hypothesized that small EVs might actively modulate biomechanical properties of target cells, key step in metastasis formation and progression. To test such assumption, we examined the effects of small EVs derived from the TNBC MDA-MB-231 breast cancer cells on the stiffness, cytoskeleton organization, nuclear/cellular morphology, and Yap activity of the Luminal A MCF7 breast cancer cells. The two cell lines have been chosen according to their different well-defined metastatic potential: MDA-MB-231 cells are characterized by high proliferation, motility, and metastatic rate (Islam & Resat, 2017), whereas MCF7 cells are tumorigenic but have low/absent metastatic potential (Gest et al., 2013; H. et al., 2011).

### 3.1 Small Extracellular Vesicle isolation and characterization

Since the experimental conditions used for the cell growth can affect the EV recovery and the final EV sample obtained, experimental procedures were fine-tuned and standardized, as much as possible, in order to maximize the number of controlled parameters for vesicle isolation (e.g. cell culture passage number, seeding confluence, and regular checks for Mycoplasma contamination) and characterization, as suggested by MISEV2018 guidelines (Thery et al., 2018).

Small EVs were isolated from MDA-MB-231 cells by ultracentrifugation (hereinafter referred to as “231_sEVs”), as described in Materials and Methods. The isolated 231_sEVs were characterized from a morphological, dimensional, and biomolecular point of view. SEM images allowed to recognize the typical rounded structure (Figure 1a) and the small EV diameter distribution, ranging from 40 to 200 nm (Figure 1b), of the isolated 231_sEVs. The spherical vesicle shape of 231_EVs was confirmed also through AFM imaging (Figure 1c); also vesicle heights and diameters (from 10 to 60 nm and 30 to 160 nm, respectively) derived from AFM analysis (Figure 1d) comply with those obtained in the literature (Parisse et al., 2017; Sharma et al., 2010).

**Figure 1.**
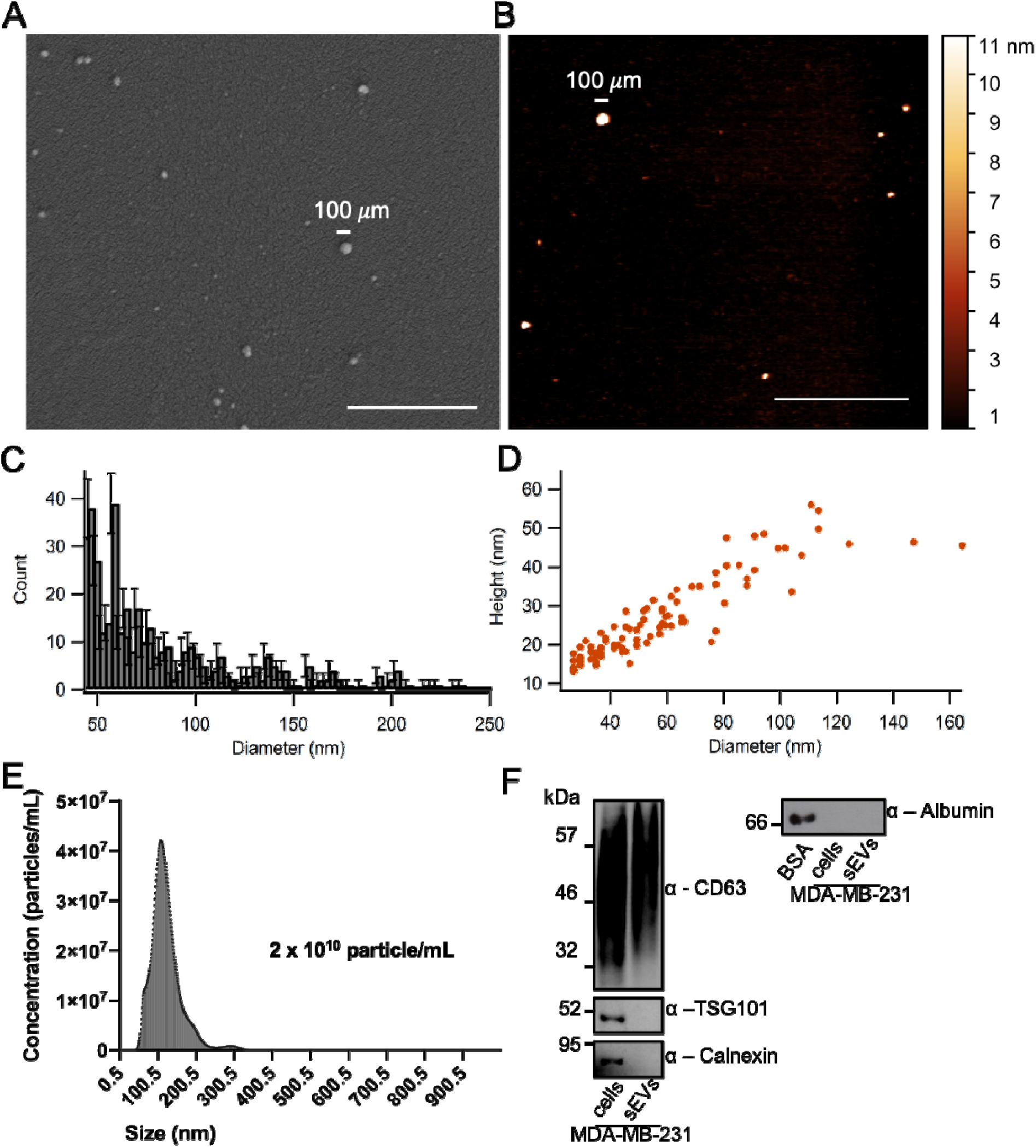
MDA-MB-231-derived small extracellular vesicles (231_sEVs) characterization. **a**) A representative SEM image of 231_sEVs. **b**) A representative AFM image of 231_sEVs. **c**) Diameter histogram of 231_sEVs obtained from the analysis of SEM images (Poisson error bars). **d**) Scatterplot of height and diameter vesicles obtained from the analysis of AFM images. **e**) Nanoparticle concentration and size distribution of 231_sEVs obtained through NTA. **f**) Western blot analysis of vesicle markers (CD63 and TSG101), and cellular (Calnexin) and serum (Albumin) contaminants in both MDA-MB-231 cellular lysate and 231_sEVs. Scale bar in **a**) and **b**) indicates 1 µm.

NTA measurements of 231_sEVs showed a vesicle concentration of 2 × 10^10^ particle/mL and a size distribution with a modal value of ∼150 nm (Figure 1e), which falls within the typical small EV diameter range (Thery et al., 2018). Western blot analysis indicated the presence of two typical small EV marker proteins: CD63 in both MDA-MB-231 cell lysate and 231_sEVs and TSG101 in cell lysate only; moreover, the analysis revealed the absence in 231_sEVs of the serum contaminant albumin and the endoplasmic reticulum-specific marker calnexin, which was detected only in MDA-MB-231 cellular lysate (Figure 1f).

### 3.2 Functional experiments: effects of MDA-MB-231-derived small Extracellular Vesicles in MCF7 cells

A quantitative estimation of 231_sEVs for functional experiments was performed by evaluating their protein concentration via Bradford assay (range between 0.6 and 1.1 µg/µL). The two breast cancer cell lines, MCF7 and MDA-MB-231, have very different properties (stiffness, cytoskeleton, nuclear/cellular morphology, and Yap activity). Therefore, we decided to optimize the acquisition parameters of the experiments separately for the two cell lines and then at a later time for the MCF7 treated with 231_sEVs and MCF7 control (addition of PBS), in order to better highlight the differences between the latter two.

#### 3.2.1 Small Extracellular Vesicles derived from MDA-MB-231 promote proliferation in MCF7 cells

Cell proliferation assay was first performed on MCF7 and MDA-MB-231 cells, confirming the higher proliferation of the latter, in agreement with the literature (Figure 2a) (Gest et al., 2013; H. et al., 2011; Islam & Resat, 2017). The activity and functionality of the isolated 231_sEVs were evaluated by verifying differences in MCF7 cell proliferation upon the addition of 231_sEVs. In order to optimize cellular treatment, different vesicle concentrations (0.05 µg/µL, 0.1 µg/µL, and 0.2 µg/µL) after 48 hours of incubation times were tested (Figure Supplementary 1). The greatest effect in proliferation was observed in the case of the highest small EV concentration, 0.2 µg/µL. This condition, in comparison with the relative control, is shown in Figure 2b. No significant or lower increase in cellular proliferation was observed for all the other conditions (Figure Supplementary 1). This result indicates that isolated 231_sEVs are active and can transfer molecular information from donor to target cells.

**Figure 2.**
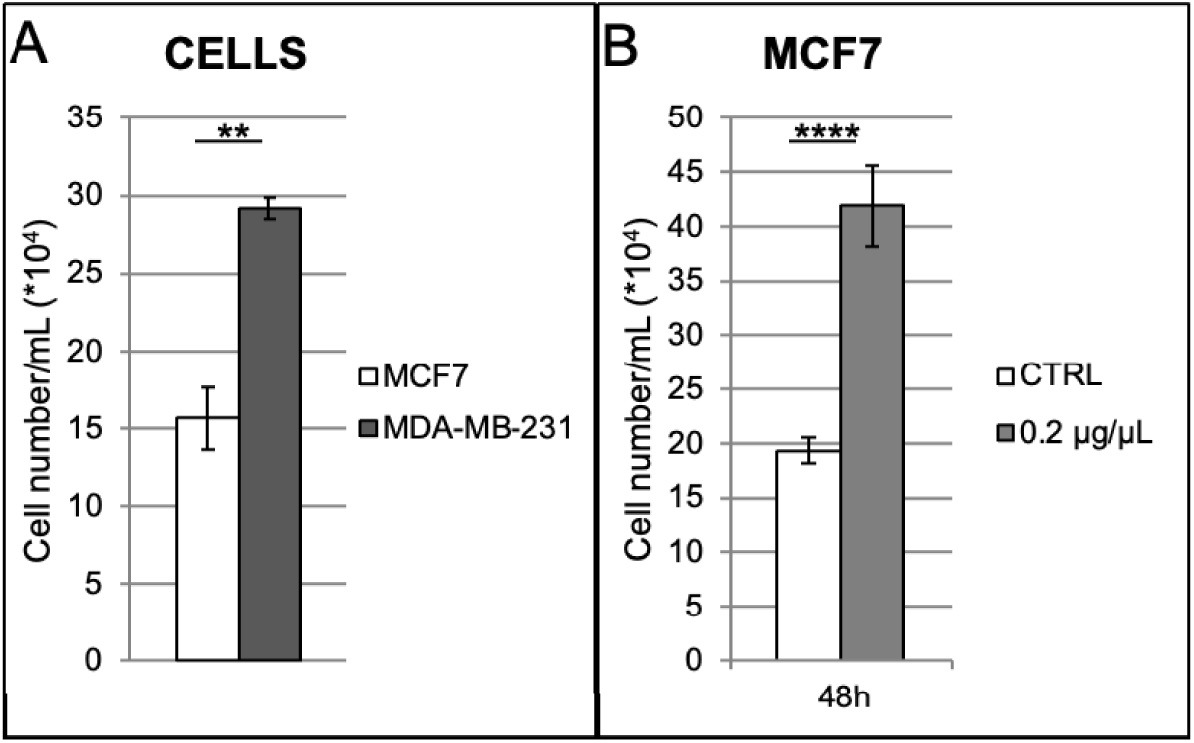
Effects of small EVs derived from MDA-MB-231 on MCF7 cell proliferation. **a**) Cell proliferation of MCF7 and MDA-MB-231 cells in comparison. **b**) Cell proliferation of MCF7 cells treated with 231_sEVs, in relation to their untreated control. Data are expressed as mean ± SD. Significance of data differences was established via two-tailed Student’s t-test.

#### 3.2.2 Small Extracellular Vesicles derived from MDA-MB-231 induce biomechanical changes in MCF7 cells

It is known from the literature that stiffness (which can be parameterized by the Young’s modulus) of single cancer cells, and in particular of most metastatic ones, is lower compared to the one of healthy cells for various cancer types (Lekka, 2016). Here, we set out to evaluate if TNBC-derived small EVs can modulate cellular stiffness of recipient cells.

MCF7 and MDA-MB-231 cell stiffness was measured by means of Force Spectroscopy AFM. One force-distance curve was acquired for each cell by indenting with a silicon spherical bead (20 µm of diameter) up to about 10 % (500 nm) of the total cell height (∼ 5 µm), previously measured via AFM morphological imaging (data not shown). The Young’s modulus can be extracted by fitting the curve within the approximation of elastic deformation (Hertz model). As expected, the MDA-MB-231 cells resulted significantly softer than the MCF7 cells (Figure 3a). When treated with 231_sEVs at the same conditions as before, the distributions of MCF7 cell stiffness values decrease with respect to the control (Figure Supplementary 2), but only MCF7 cells treated with 0.2 μg/μL of 231_sEVs gave a significant decrease, reported in Figure 3b; two representative force-displacement curves of both MCF7 and MCF7 treated with 231_sEVs were shown in Figure 3c. To be sure that such effect is related to the specific molecular content of the 231_sEVs and not to the addition of any EVs, MCF7 cells were treated with 0.2 μg/μL of small EVs isolated from the MCF7 cells themselves (referred to as “7_sEVs”), as control. As shown in Figure Supplementary 3, no significant difference can be observed between the stiffness of MCF7 cells treated with 7_sEVs and their relative control.

**Figure 3.**
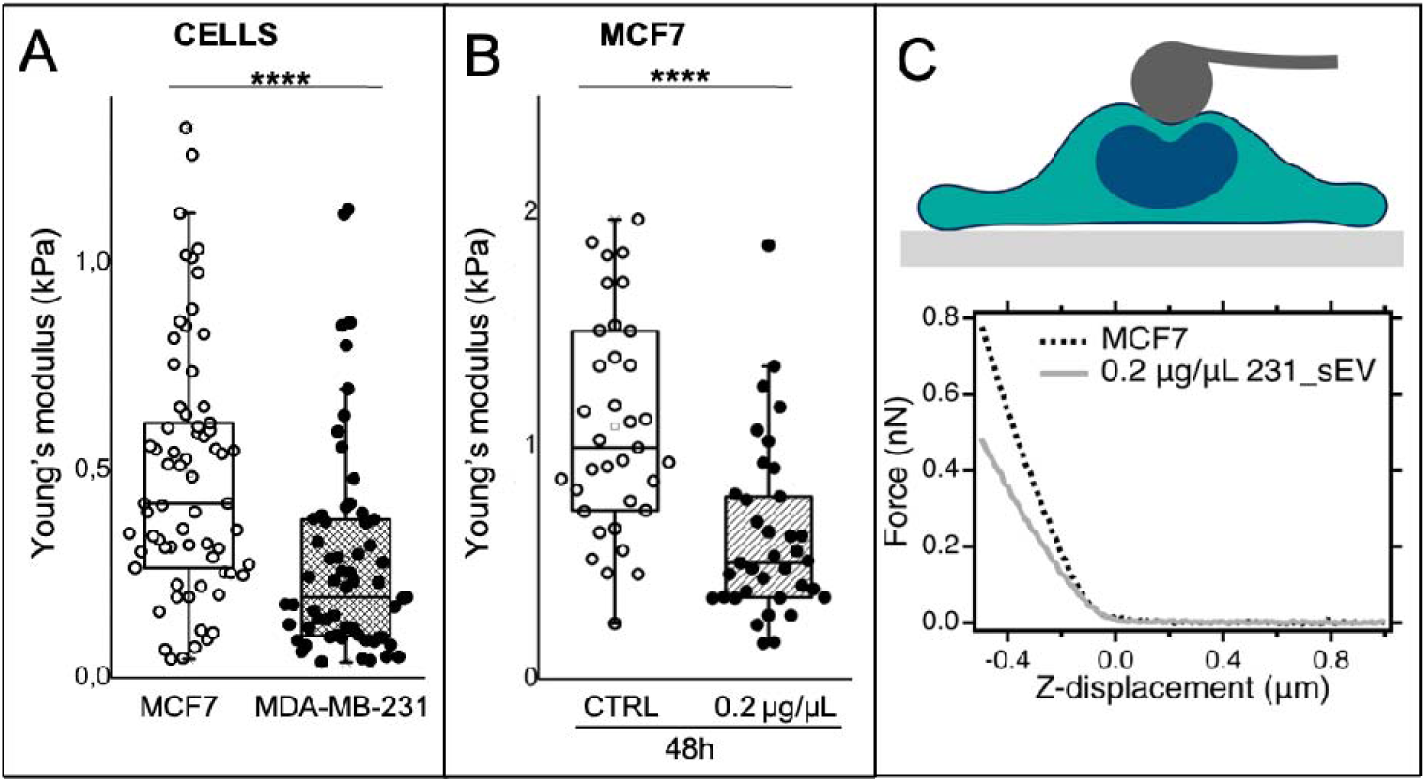
Effects of small EVs derived from MDA-MB-231 on the MCF7 cell stiffness. **a**) Boxplot showing the Young’s modulus distributions of MCF7 and MDA-MB-231 single cells and **b**) of MCF7 cells after the addition of 231_sEVs (0.2 µg/µl for 48 hours), in relation to their negative control. The lower and the upper boundaries of the box represent Q1 (25 percentile) and Q3 (75 percentile) of the data, respectively; the ▫ symbol and the horizontal bar inside the box represent the mean and median, respectively. Significance of data differences was established via Wilcoxon test. **c**) Representative AFM force-displacement curve of MCF7 cells and MCF7 cells upon the 231_sEV addition.

Therefore, these results suggest that MDA-MB-231-derived small EVs can transfer molecular information to MCF7 target cells that is responsible for the acquisition of a biomechanical phenotype similar to donor cells.

#### 3.2.3 Small Extracellular Vesicles derived from MDA-MB-231 induce cytoskeleton and nuclear/cellular morphology rearrangements in MCF7 cells

For a comprehensive understanding of the observed biomechanical changes, we explored cytoskeleton, adhesion, and nuclear/cellular morphology of MCF7 cells treated or not with 231_sEVs. Indeed, cytoskeleton organization is often associated with malignant transformation (Alibert et al., 2017) and it has important roles in cellular biomechanical modulation (Fritsch et al., 2010). Actin organization and focal adhesions (FAs) appear to play a major role in the regulation of cellular biomechanics (Tavares et al., 2017). Moreover, in cancer cells, morphology and stiffness of the nucleus, the most rigid cellular organelle, are altered when compared with healthy cells (Chiotaki et al., 2014)(Fischer et al., 2020). In particular, nuclear irregularity, deformity and softening, as result of alteration of the nucleoskeleton and nucleus-cytoskeleton interactions, are associated to high tumoral invasiveness of cancer cells (Chiotaki et al., 2014)(Senigagliesi et al., 2019). Cytoskeleton and nuclear properties of MCF7 and MDA-MB-231 breast cancer cells have already been extensively analyzed (Chiotaki et al., 2014), and turned out to be very different.

Here, significant differences were observed in F-actin (stress fibers and cortical) by comparing both the MCF7 to the MDA-MB-231 cells and the MCF7 cells treated with 231_sEVs to their control (Figure 4a-b). F-actin fluorescence intensity in both MDA-MB-231 cells and MCF7 cells after the addition of 231_sEVs resulted higher compared with the MCF7 cells and MCF7 treated with PBS (control), respectively, as shown in Figure 4 (n = 240 MDA-MB-231 cells and n = 443 MCF7 cells; n = 790 MCF7 CTRL cells and n = 849 231_sEVs-treated MCF7 cells). No differences were observed in cell area of all samples taken into consideration (Figure Supplementary 4). Phalloidin staining was used to assess cell shape, more specifically the elongation factor of each cell, as indicator of the spindle-like morphology of mesenchymal-like cells. The elongation factor *E* is defined as E = A_L_/A_S_ – 1, where A_L_ and A_S_ represent the long and short axis of the cell, respectively. *E* corresponds to zero for a circle, and one for an ellipse with an axis ratio 1:2. Cells that presented *E* values 0.0 – 0.5 were considered as spherical, while values equal or higher than 0.5 as elongated (Stylianou et al., 2018). MDA-MB-231 cells have the typical elongated mesenchymal-like morphology compared to the MCF7 cells, as shown in epifluorescence images and relative histograms of Figure 5a (n = 24 MDA-MB-231 cells and n = 28 MCF7 cells).

**Figure 4.**
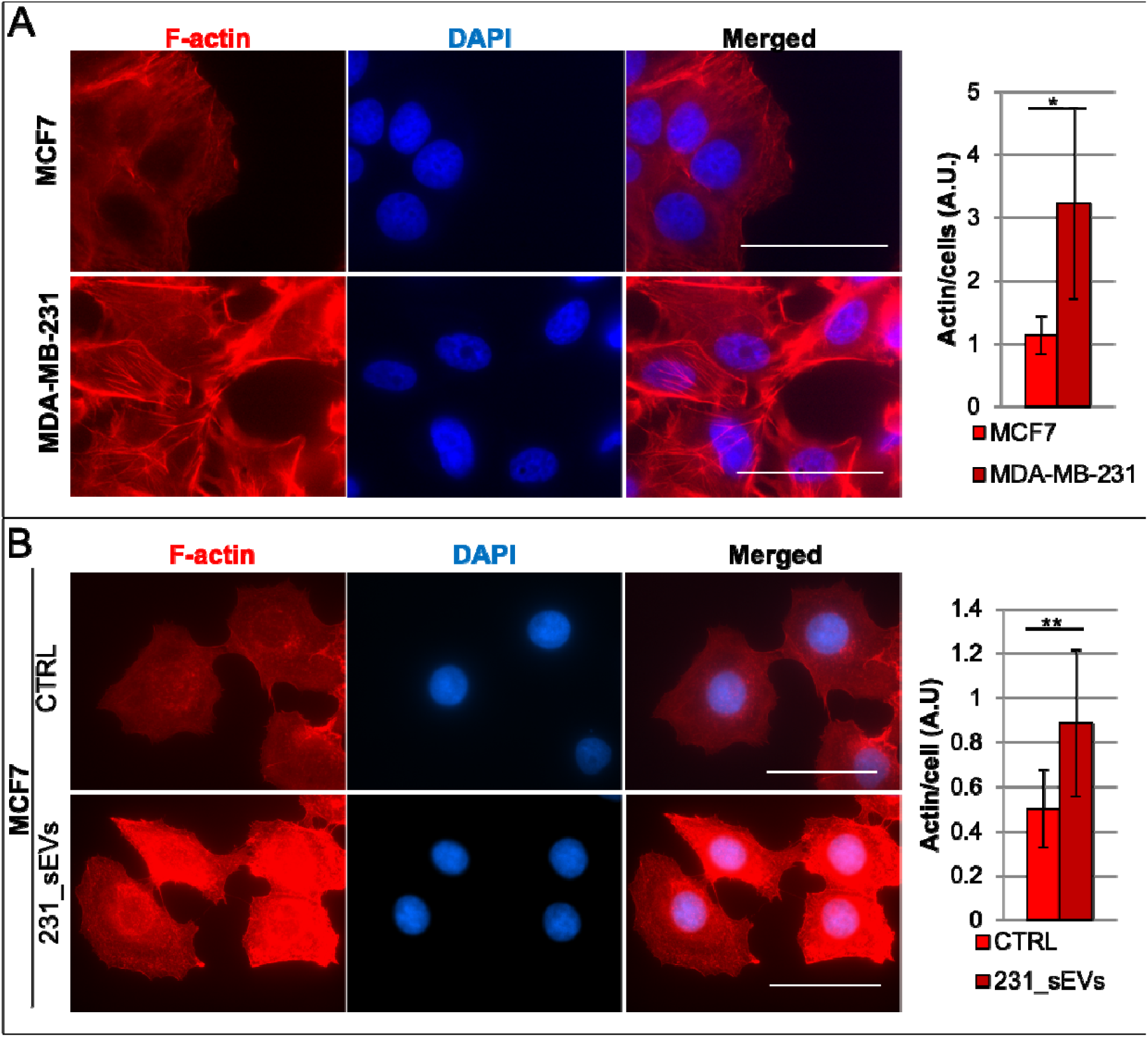
Effects of MDA-MB-231-derived small EVs on F-actin of MCF7 cells. Representative epifluorescence images on left and relative histograms on right, respectively, showing the F-actin of **a**) MCF7 and MDA-MB-231 cells and of **b**) MCF7 after the addition of 231_sEVs, in relation to their relative control. Data are expressed as mean ± SD. Significance of data differences was established via two-tailed Student’s t-test. Scale bar indicates 50 µm.

**Figure 5.**
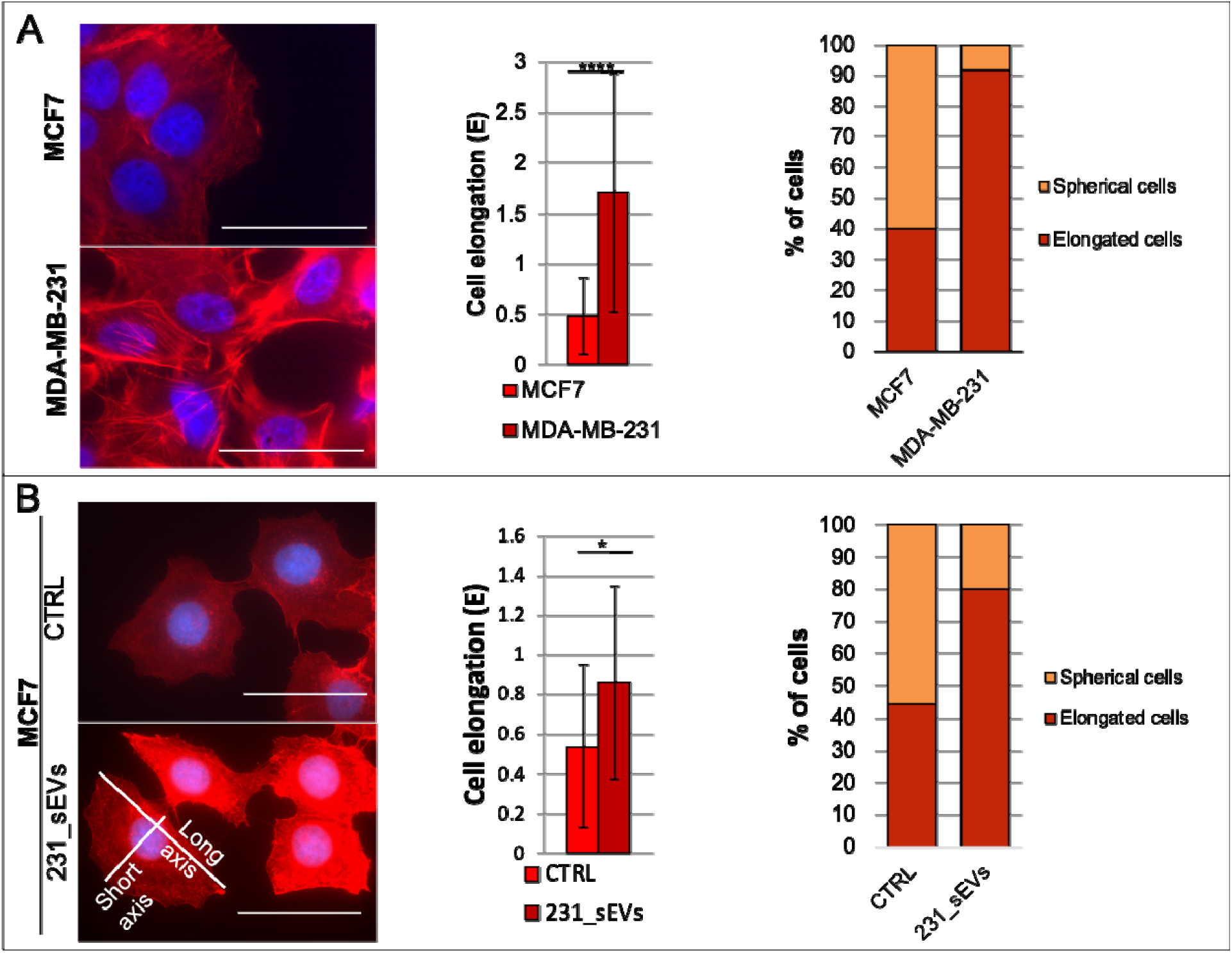
Effects of 231_sEVs on cell elongation of MCF7 cells. Representative epifluorescence images on left and histograms on right, respectively, showing the cell elongation and percentage of elongated cells in (**a**) MCF7 and MDA-MB-231 cells and in (**b**) MCF7 cells treated with 231_sEVs, in relation to their relative control. Data are expressed as mean ± SD. Significance of data differences was established via two-tailed Student’s t-test. Scale bar indicates 50 µm.

When treated with 231_sEVs, MCF7 cells showed a substantial increase in the percentage of elongated cells, compared to their control, as shown in Figure 5b (n = 25 MCF7 CTRL cells and n = 25 231_sEVs-treated MCF7 cells). This rearrangement in shape makes the MCF7 cell morphology similar to that of MDA-MB-231 cells and strongly suggests an epithelial-mesenchymal transition phenomenon, due to the 231_sEV activity.

To further investigate the small EV-induced transformations, we also investigated the nuclear morphology of the different cells. Significant smaller nuclear area and lower nuclear circularity were observed in MDA-MB-231 with respect to MCF7 cells, as shown in Figure 6a (n = 525 MDA-MB-231 cells and n = 744 MCF7 cells). Similarly, MCF7 cells incubated with 231_sEVs show a significant reduction in nuclear size and circularity, when compared with the control (Figure 6b; n = 744 MCF7 CTRL cells and n = 712 231_sEVs-treated MCF7 cells). Therefore, the activity of vesicles derived from MDA-MB-231 makes nuclear morphology of the MCF7 cells similar to that of MDA-MB-231 cells.

**Figure 6.**
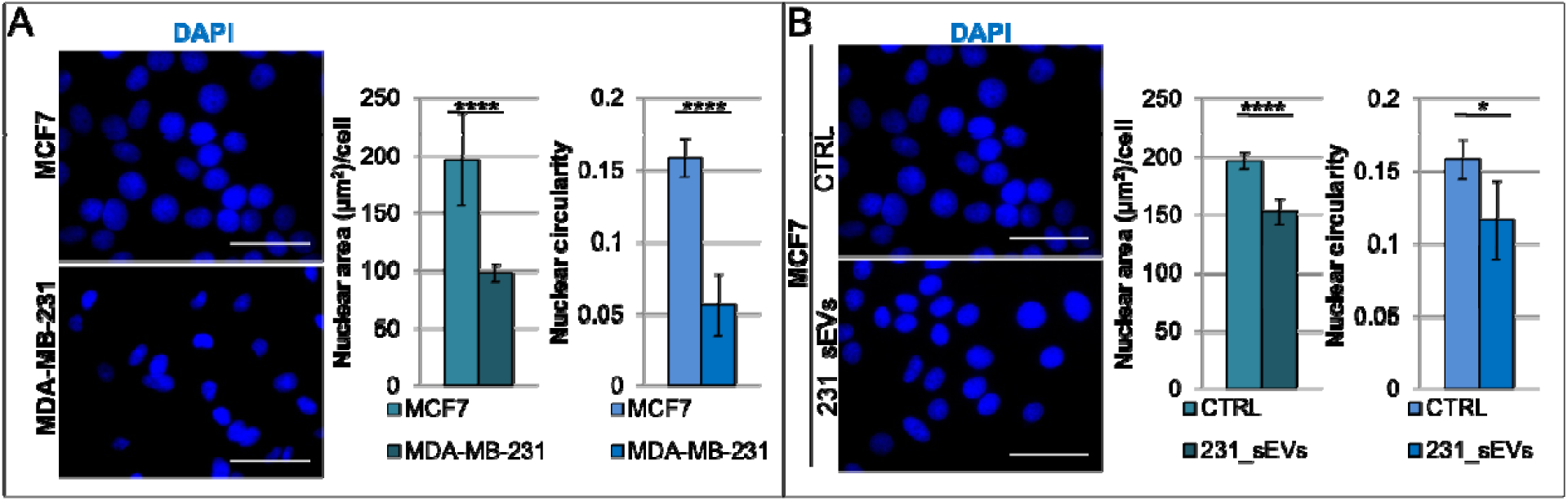
Effects of 231_sEVs on nuclear morphology of MCF7 cells. Representative epifluorescence images on left and histograms on right, respectively, showing the nuclear area and nuclear circularity in (**a**) MCF7 and MDA-MB-231 cells and in (**b**) MCF7 cells treated with 231_sEVs, in relation to their relative control. Data are expressed as mean ± SD. Significance of data differences was established via two-tailed Student’s t-test. Scale bar indicates 50 µm.

Lastly, we examinated both the density and size of FAs of the different cells by staining the Vinculin and the phosphorylated-focal adhesion kinase (pFAK).

When compared with MCF7, MDA-MB-231 cells show significantly lower density of FAs, as inferred from both the Vinculin and pFAK TIRF images, and significant lower adhesion size, as detected via the pFAK signal (Figure 7a-b; n = 47 and n = 35 MDA-MB-231 cells for Vinculin and pFAK, respectively; n = 25 and n = 29 MCF7 cells for Vinculin and pFAK, respectively). On the contrary, MCF7 cells treated with 231_sEV show a significantly higher density and size of focal adhesions, when compared to the control (Figure 7c-d; n = 25 and n = 29 MCF7 CTRL cells for Vinculin and pFAK, respectively; n = 26 and n = 23 231_sEVs-treated MCF7 cells for Vinculin and pFAK, respectively). Therefore, in contrast to the cytoskeleton and morphology results (Figure 4-5), 231_sEVs appear to provide MCF7 cells an adhesion phenotype different from the donor MDA-MB-231 cells. Specifically, they promote an increase of FA activity.

**Figure 7.**
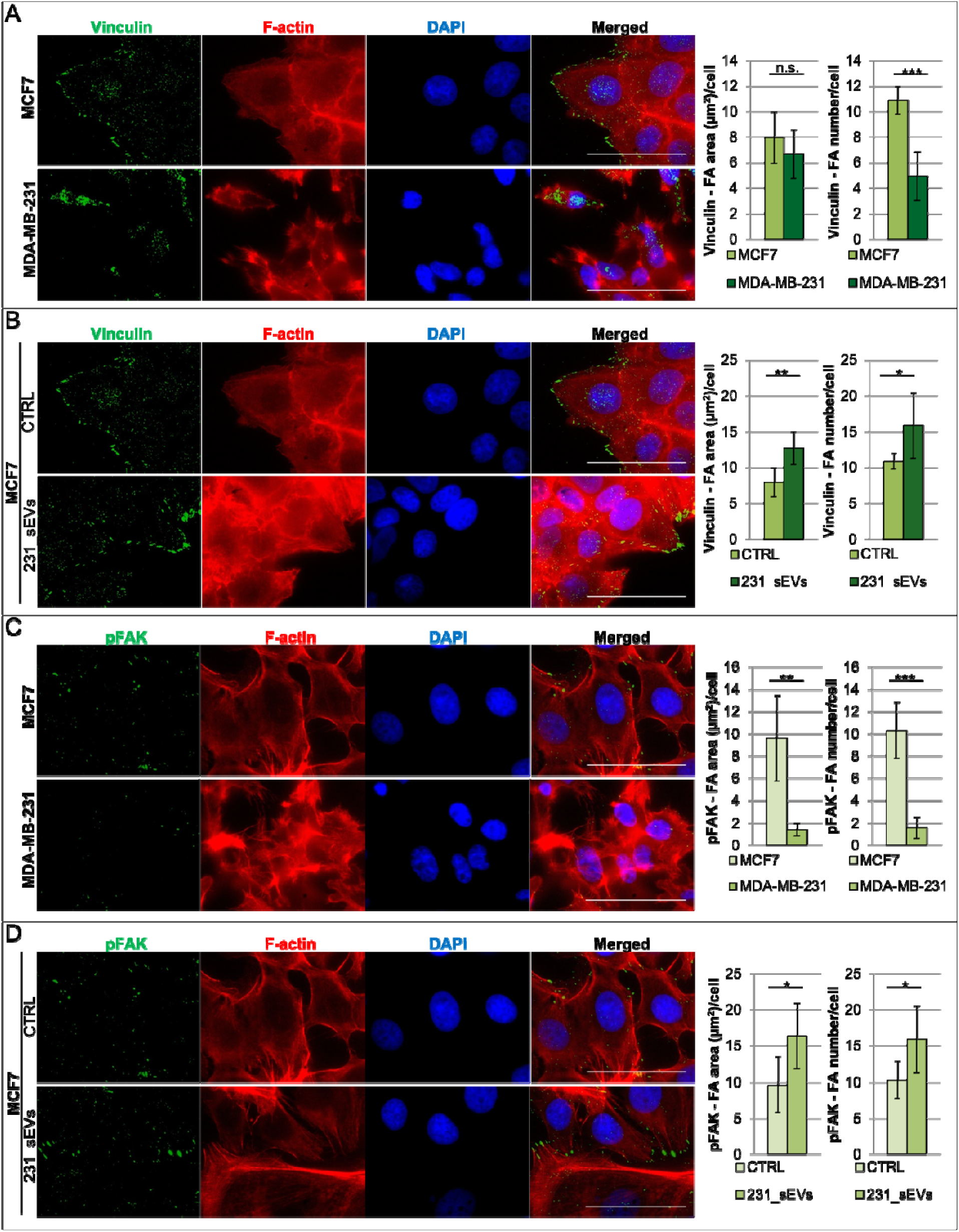
Effects of small EVs derived from MDA-MB-231 on focal adhesions of MCF7 cells. Representative TIRF (Vinculin and pFAK) and epifluorescence (F-actin and DAPI) images on left and relative histograms on right, respectively, showing the area, density, and activity of FAs in (**a-c**) MCF7 and MDA-MB-231 cells and in (**b-d**) MCF7 cells treated with 231_sEVs, in relation to their negative control. Data are expressed as mean ± SD. Significance of data differences was established via two-tailed Student’s t-test. Scale bar indicates 50 μm. N.s. indicates not significant.

#### 3.2.4 Small Extracellular Vesicles derived from MDA-MB-231 induce Yap activation in MCF7 cells

We then wondered if the phenotypic rearrangement previously observed could match with any variations in Yap activity. Yes-associated protein (YAP), a transcriptional co-activator negatively regulated from the Hippo pathway, is known to play an important role in cellular biomechanical modulation and, in particular, in regulation of the acto-myosin network (Qiao et al., 2017)(Dobrokhotov et al., 2018) and focal adhesions (Nardone et al., 2017). Yap is an oncoprotein and abnormally accumulation of nuclear YAP has been observed in many types of cancer, including breast cancer (Qiao et al., 2017)(Dobrokhotov et al., 2018). In particular, the increase in Yap activity can drive (i) the lack of contact/density-dependent inhibition of growth (Dasgupta & McCollum, 2019), (ii) an increase in cellular motility, invasion and metastasis (Z. Wang et al., 2014), (iii) a marked increase in FA formation (Nardone et al., 2017)(Shen et al., 2018), and (iv) the promotion of FAK phosphorylation (Shen et al., 2018).

Therefore, we investigated via RT-qPCR the expression level of three Yap downstream genes, CTGF, CYR61, and ANKRD1, as a readout of Yap activation/activity. In agreement with literature (Shen et al., 2018), we found that Yap is minimally active in MCF7 cells, whereas Yap downstream genes are overexpressed in MDA-MB-231 cells (Figure 8a). Interestingly, a not significant and a significant increase in CTGF and ANKRD1 expression, respectively, was reproducibly observed in MCF7 cells upon 231_sEV treatment (Figure 8b). This result could suggest a correlation between the 231_sEV addition and an enhanced Yap activity in MCF7 cells; these variations in Yap activity, which deserves further analysis, suggest the involvement of a pathway regulated and regulator of biomechanics as proof of the cell phenotypic changes observed above in MCF7 cells treated with 231_sEVs.

**Figure 8.**
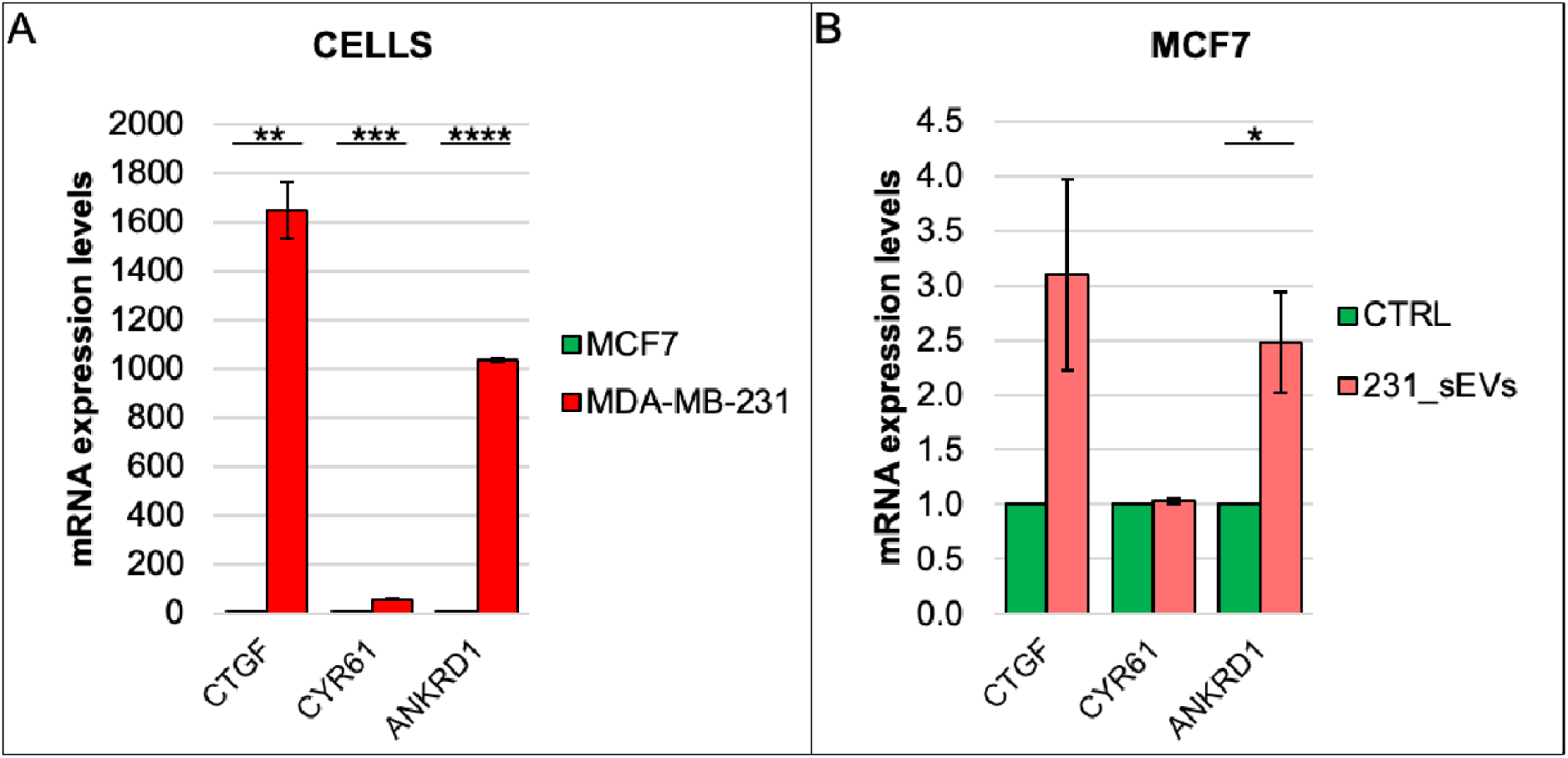
Effects of small EVs derived from MDA-MB-231 on Yap downstream gene expression of MCF7 cells. Expression of Yap target genes measured by RT-qPCR in **a**) MCF7 and MDA-MB-231 cells and in **b**) MCF7 cells treated with 231_sEVs, in relation to their negative control. Data normalized on histone H3. Data are expressed as mean ± SD. Significance of data differences was established via two-tailed Student’s t-test. N.s. indicates not significant.

## 4 DISCUSSION

Despite the constant progress in breast cancer field, the development of metastases in TNBC remains a highly complex and poorly understood process with a relatively poor outcome (Al-Mahmood et al., 2018). It is known that more deformable and softer cancer cells exhibit greater metastatic ability (Rudzka et al., 2019). On the other side, extracellular vesicles released by triple-negative breast cancer cells have been found to transfer functional cargos to target cells promoting cell proliferation, tumor growth, cancer cell invasion, and metastasis (Green et al., 2015). In light of that, we were interested in highlighting the mechanisms through which EVs from TNBC induce or regulate the metastatic processes, and in particular how do they influence/modulate the biomechanics of the recipient cells. Sparse information in this direction is coming from the recent literature: Sung and colleagues demonstrated that the secretion of small extracellular vesicles is required for directionally persistent and efficient in vivo movement of cancer cells (Sung et al., 2015); small EVs secreted by mesenchymal stromal/stem cell-derived adipocytes can promote breast cancer cell growth by activating Hippo signaling pathway (S. Wang et al., 2019); breast cancer-derived EVs were seen to contribute to metastasis by altering the tissue mechanics of distant organs to support tumor cell invasion and seeding (Pokharel et al., 2016)(Barenholz-Cohen et al., 2020); moreover, single EV characterization studies reported the conservation of biomechanical traits of EVs with respect to their cells of origin (softer EVs were found to be secreted by softer cells) (LeClaire et al., 2021; Ye et al., 2021). In the present study we showed for the first time that beside preserving the biophysical properties of the donor cells, as shown by (LeClaire et al., 2021 Ye et al., 2021), the extracellular vesicles are able to transfer these features directly to the target cells. In detail, we demonstrated that TNBC-derived small EVs directly modify the biomechanical phenotype of the non-invasive MCF7 target cells, by affecting cellular stiffness, cytoskeleton, nuclear and cellular morphology, adhesion, and Yap downstream gene expression variations. Our results evidenced that the MCF7 cells treated with 231_sEVs adapt their stiffness, cellular and nuclear morphology in the likeness of the MDA-MB-231 cells. Yet, their adhesion properties are different. Interestingly, we observed an increase in FA number, density, and activity in 231_sEVs treated MCF7 cells with respect to the EVs progenitor cells, which might be a direct consequence of small EV uptake. It has been shown from recent studies that the integrin beta 3, ITGB3, which plays an essential role in cancer metastasis in MDA-MB-231, and its interaction and activation of focal adhesion kinase (FAK has a fundamental role in extracellular vesicle biogenesis and uptake in breast cancer cells (Altei et al., 2020)(Fuentes et al., 2020). According to these works, ITGB3 is required both as small EVs receptor (by interacting with heparan sulfate proteoglycans (HSPGs)) and for intracellular FAK activation to promote vesicles endocytosis. These findings are well fitting with our observations. Although FAs and FAK in breast malignancy metastasis have been widely studied, it is not perfectly clear yet how FAs are regulated in tumors and many data present in the literature on FAs size, number, and activity in cancer cell lines are quite scattered. Direct correlation between metastatic/aggressive potential of cancer cells and large number/small size of focal adhesion complexes were demonstrated in some other works (Rönnlund et al., 2013)(Gad et al., 2012)(Kraning-Rush et al., 2012). In contrast, other studies showed that invasive cells are characterized by large dynamic adhesion sites with an increased activation of the phosphorylated-focal adhesion kinase (pFAK) (Peschetola et al., 2013)(Tavares et al., 2017). pFAK, by recruiting other signaling molecules, promotes in turn the assembly of focal adhesion complexes (Shen et al., 2018). This protein has been observed to have an oncogenic role in many types of human cancers (Shen et al., 2018), including metastasis and poor prognosis in breast cancer (M. Luo & Guan, 2010). Our FA results are consistent with the work of (Peschetola et al., 2013)(Tavares et al., 2017), pointing to a correlation of invasiveness with large dynamic adhesion sites and an increased activation of pFAK.

Moreover, we also highlighted a high expression of Yap downstream genes in the 231_sEVs treated MCF7 cells, similar to the case of MDA-MB-231 cells, and its correlation with high levels of filamentous actin, as well as with a marked increase in FA formation/promotion of FAK phosphorylation. Also in the case of the oncoprotein Yap, conflicting results on the mechanisms through which it regulates actin cytoskeleton polymerization and FAs were reported. Concerning the actin regulation, in the study of Qiao et al. the Yap activation in human gastric cancer cells is shown to drive an actin depolymerization with consequent cell softening (Qiao et al., 2017), while in the study of Nardone et al. it promotes in adipose tissue-derived mesenchymal stem cells the formation of stress fibers with consequent cell stiffening (Nardone et al., 2017). Such opposite results might be related to the way Yap regulates the acto-myosin system, as it promotes the expression of both a RhoA activator (ARHGEF17) and RhoA inhibitors (ARHGAP18/29) (Dobrokhotov et al., 2018). Concerning FAs modulation, Shen et al. observed that YAP promotes focal adhesion dynamics in breast cancer cells (MCF7 and MDA-MB-231), but they did not report any significant actin changes upon Yap overexpression or silencing (Shen et al., 2018). Our findings support both the results of Nardone et al. and Shen et al. (Nardone et al., 2017)(Shen et al., 2018).

In conclusion, we showed that TNBC-derived small EVs alter the biomechanical response of recipient cells. Our results admit two possible explanations of the mechanism through which TNBC-derived small EVs can influence cell biomechanics. The first possible route is based on the hypothesis that the small EV uptake occurs through FAs, which being subject to changes, then, consequently modulate Yap activity, cytoskeleton, nuclear/cell morphology, and biomechanics of target cells. Alternatively, TNBC-derived small EVs could directly cause an increase in Yap activity, which in turn leads to cytoskeleton, nuclear/cell morphology, adhesion and, finally, biomechanical rearrangements in MCF7 cells. Further studies (as proteomic and/or RNA analyses) will be needed to clarify this point, which falls outside the aim of the present work.

We believe that testing the biomechanical properties of both cells and tissues after EV treatment might represent a new approach capable of assessing the activity of EVs and the cargo release mechanisms; this assay could be potentially applied not only to other types of cancers but also to other diseases, where biomechanical properties can have prominent roles.

## ACKNOWLEDGMENTS

The authors wish to thank the Structural Biology Laboratory of Elettra-Sincrotrone Trieste S.C.p.A for the instrumentation and the continuous support. The work was supported by Università degli Studi di Trieste, Area Science Park, European Regional Development Fund and Interreg V-A Italia - Austria 2014-2020 (EXOTHERA-ITAT1036), AIRC (IG 21803 to L.Co.) and by Regione Friuli Venezia Giulia (legge regionale 17/2004, BioMec project).

