## supplemental figures for "Triple Negative Breast Cancer-derived Small Extracellular Vesicles as Modulator of Biomechanics in target cells"

**Short title: small EVs as modulator of cell biomechanics**

Beatrice Senigagliesi ^1,2^, Giuseppe Samperi ^3^, Nicola Cefarin ^4^, Luciana Gneo ^5^, Sara Petrosino ^6^, Mattia Apollonio ^7^, Federica Caponnetto ^8^, Riccardo Sgarra ^7^, Licio Collavin ^7^, Daniela Cesselli ^8^, Loredana Casalis ^2^*, Pietro Parisse ^2,4^*

*1 Scuola Internazionale Superiore di Studi Avanzati, 34127 Trieste, Italy;;*

*2 Elettra-Sincrotrone Trieste S.C.p.A., Trieste, Italy;;*

*3 University of Parma;*

*4 Istituto Officina dei Materiali Consiglio Nazionale delle Ricerche, TASC, Trieste, Italy;;*

*5 University of Oxford; Oxford OXI 2JD, GB;;*

*6 Telethon Institute of Genetics and Medicine, Naples, Italy;;*

*7 Department of Life Science, University of Trieste, 34127 Trieste, Italy;;*

*8 Pathology Department, University Hospital of Udine, P.le S. Maria della Misericordia, 33100 Udine (UD), Italy;;*

**Supplementary Figures**

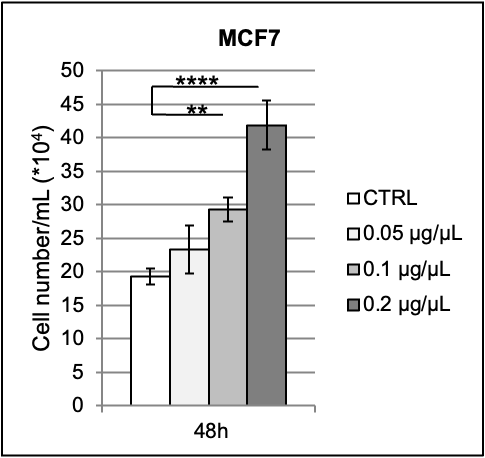

**Figure Supplementary 1. Effects of small EVs derived from MDA-MB-231 on MCF7 cell proliferation.** Cell proliferation of MCF7 after the addition of 231_sEVs at different conditions, in relation to their negative control. Data are expressed as mean ± SD. Significance of data differences was established via one-way Anova test.

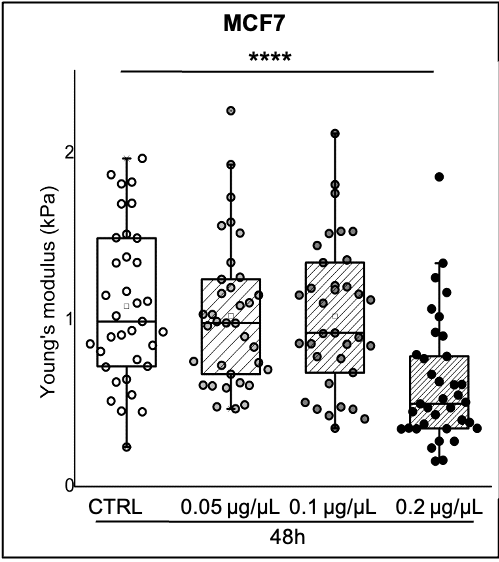

**Figure Supplementary 2. Effects of small EVs derived from MDA-MB-231 on the MCF7 cell stiffness.** Boxplot showing the Young’s modulus distributions of MCF7 after the addition of 231_sEVs at different conditions, in relation to their negative control. The lower and the upper boundaries of the box represent Q1 (25 percentile) and Q3 (75 percentile) of the data, respectively; the ▫ symbol and the horizontal bar inside the box represent the mean and median, respectively. Significance of data differences was established via Kurskal-Wallis one-way Anova test.

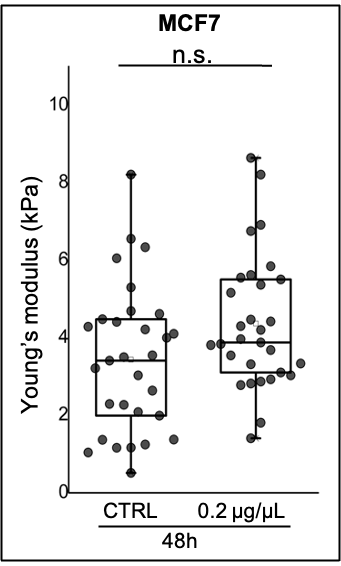

**Figure Supplementary 3. Effects of small EVs derived from MCF7 on the MCF7 cell stiffness.** Boxplot showing the Young’s modulus distributions of MCF7 after the addition of 7_sEVs at different conditions, in relation to their negative control. The lower and the upper boundaries of the box represent Q1 (25 percentile) and Q3 (75 percentile) of the data, respectively; the ▫ symbol and the horizontal bar inside the box represent the mean and median, respectively. Significance of data differences was established via Wilcoxon test. N.s. indicates not significant.

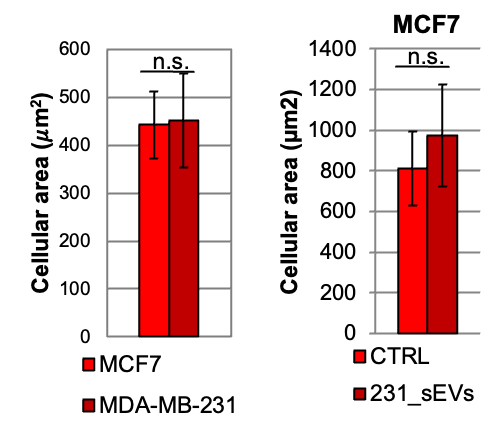

**Figure Supplementary 4. Effects of 231_sEVs on cell area of MCF7 cells.** Histograms obtained from epifluorescence images showing the cell area of cells in MCF7 and MDA-MB-231 cells and in MCF7 cells treated with 231_sEVs, in relation to their relative control. Data are expressed as mean ± SD. Significance of data differences was established via two-tailed Student’s t-test. Scale bar indicates 50 µm.
